# Humans exhibit demographic buffering against variability and persistence in inflation environments

**DOI:** 10.64898/2026.06.03.729782

**Authors:** Rahul Mondal, Samuel JL Gascoigne, Udaya S Mishra, Roberto Salguero-Gomez

## Abstract

Inflation affects human fertility and survival; persistent inflation can translate into fluctuations in annual population growth rate. Existing studies have yet to systematically examine inflation’s impact on long-term growth rates across populations. Here, we examine whether high inflation variance and autocorrelation amplify long-term growth rate’s sensitivity to inflation, across 32 countries over 1971-2024. Additionally, we examine the effect of increasing inflation variance and autocorrelation on humans’ capacity for demographic buffering, *i.e*., reducing the temporal variance in vital rates most sensitive to growth. First, we construct inflation-dependent matrix population models using a hierarchical Bayesian framework. We then calculate the stochastic sensitivity and elasticity of population growth rate to inflation to quantify demographic buffering across a variance-autocorrelation parameter space. We find that the stochastic sensitivities of the growth rate increase with inflation’s variance and autocorrelation. However, we do not find support for the expected adverse effects of inflation variance and autocorrelation on humans’ capacity to buffer against inflation. Our framework provides a valuable tool for examining the impact of resource uncertainty on life history strategies in stochastic environments. We discuss how our framework can be extended to study the resource-dependent population dynamics within variable environments and across the tree of life.

## Introduction

Uncertainty in inflation rates can lead to significant variation in national fertility rates. Empirical evidence suggests that higher inflation uncertainty leads to increased uncertainty in income and precautionary savings (Fischer et al. 2024), leading to a reduction in both planned household spending and fertility (Gozgor et al. 2021; Ranjan 1999). Moreover, individuals often postpone pregnancy during periods of economic uncertainty (Sobotka et al. 2011), particularly pregnancy for the first child (Kreyenfeld and Andersson 2014). Specifically, perceived economic uncertainty, resulting from escalating global changes in technology, media, migration, and climate, is suggested to play an increasingly significant role in shaping fertility decisions (Hellstrand et al. 2024; Vignoli et al. 2020). For example, a recent Finnish survey found that perceived uncertainty was one of the primary reasons for fertility postponement in the aftermath of the 2010s (Savelieva et al. 2023). Additionally, evidence of fertility responding to economic uncertainty has been reported across Latin America (Adsera and Menendez 2011) and Europe (Vignoli et al. 2020). In turn, inflation may be a driver linking economic uncertainty and fertility, as higher realised inflation can lead to elevated inflation uncertainty, income uncertainty, a household’s perceived risk of unemployment (Fischer et al. 2024) and thereby an adverse effect on fertility.

Inflation uncertainty can also influence human survival through precautionary savings, reducing the demand for health. However, literature exploring the inflation-survival nexus is scarce. There is empirical evidence suggesting that humans cope with high inflation environments by reducing expenditure on food, purchasing lower-quality food, or omitting an entire meal (Fouéré et al. 2000). Furthermore, studies conducted in Sub-Saharan Africa indicate that children may be particularly vulnerable to malnutrition and associated physical growth constraints due to inflation (Delpeuch et al. 1996). Interestingly, a study by Williams et al. (Williams et al. 2016) on 21 Latin American countries found that inflation is associated with increased child and male adult mortality. Moreover, studies examining the relationship between a high inflation environment and suicide rates suggest a potential positive association (Andreeva et al. 2008). Notwithstanding the growing literature on inflation and population health, the relationship between inflation and long-term population through survival rates remains largely unexplored.

Examining the nexus among the economy and fertility/survival has become one of the most active areas of human demographic research. Existing studies suggest that multidimensional factors, including rising wages and female labour force participation, changing social norms about marriage, and family structures amid rising living costs, have driven a global trend of fertility decline (Bloom et al. 2024; Doepke 2004). Additionally, the quality-quantity trade-off, *i.e*., having many children versus spending more resources per child, gradually shifted towards quality with a reduction in child mortality (Barro and Becker 1989; Doepke 2004). Among populations in the “lowest-low-fertility regime”, such as Finland, Italy, Norway, and Spain, economic uncertainty is argued to be a game-changer in driving the fertility dynamics (Kohler et al. 2002; Vignoli et al. 2020). On the other hand, the impact of economic uncertainty on population health and mortality is a burgeoning area of research (Abdou et al. 2022; Kalemli-Ozcan and Weil 2010; Kawachi et al. 2023). However, the conflating mechanisms associated with education, the socio-political environment, and the aforementioned global changes make it difficult to isolate the effect of economic uncertainty on fertility or survival (Hellstrand et al. 2024; Matysiak et al. 2021; Vignoli et al. 2020). Furthermore, to our best knowledge, no studies have yet examined the sensitivity of population growth to the marginal effect of economic uncertainty on human fertility and survival. Therefore, in this study, we examine the sensitivity of the long-term stochastic population growth rate to the marginal effect of economic uncertainty, operationalised here by inflation uncertainty, on the demographic rates.

Understanding the long-term effect of inflation on vital rates is of profound importance as inflation can influence asymptotic population dynamics. Hitherto, most studies examining the relationship between inflation or economic uncertainty and human demographic behaviour have spanned over relatively short-term periods (Adsera and Menendez 2011; Comolli 2017), and been regional in nature (Vignoli et al. 2020; Williams et al. 2016). However, results from short-term regional studies limit inferences about the influence of inflation uncertainty on human life history strategies, as they are sensitive to transient inflationary trends/shocks specific to each region (*e.g*., Adsera and Menendez, 2011). Interestingly, increased uncertainty about one-year-ahead inflation significantly raises uncertainty about five-year-ahead inflation, and short-run inflation expectations drive long-run inflation expectations (Fischer et al. 2024). Therefore, long-term inflation uncertainty may impose selective pressure on survival and fertility, and consequently life history strategies. An evolutionary framework that examines the responses of vital rates within the context of adaptive optimisation under a variable inflation environment is essential for understanding the effect of inflation on long-term population dynamics (Caswell 2001; Tuljapurkar 1982a, 1982b).

The demographic buffering hypothesis (DBH, hereafter, Pfister 1998) provides the essential framework to understand the relationship between long-term population dynamics and its environmental drivers (Hilde et al. 2020) - inflation in the case of human populations. Specifically, the DBH predicts a negative correlation between the sensitivity of population growth rate to its underlying vital rates and the temporal variation of said vital rates (Hilde et al. 2020; W. F. Morris and Doak 2004). Therefore, examining how an environmental driver, such as inflation, affects the capacity of a population to reduce the temporal variance of vital rates can be used to understand the influence of the driver on life-history strategies and long-term population dynamics. The biodemographic literature quantifies demographic buffering as the sum of the elasticity of stochastic population growth to the variance of vital rates 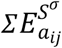 (Santos et al. 2024), and examines how 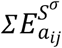 depends on the variance and temporal autocorrelation of the environmental variable of interest (Gascoigne et al. 2025; Haridas and Tuljapurkar 2005; Tuljapurkar and Haridas 2006). Hence, we can use this demographic buffering framework to quantify the effect of variability in inflation on long-term human population growth via its impact on vital rates.

Here, we investigate the impact of variability in inflation rates on the ability of human populations to remain demographically buffered, that is for the population growth rates to remain constant regardless of the variability in key environmental drivers. To explore this question, we utilise high-resolution inflation and demographic data to construct inflation-dependent Bayesian matrix population models (MPM) (Wheldon et al. 2015; Wiśniowski et al. 2015). Specifically, we use these MPMs to test the following hypotheses: (H1) With increasing variance and temporal autocorrelation in inflation rates, the long-term stochastic population growth rate *λ*_*s*_ becomes increasingly sensitive to the effect of inflation on fertility. The rationale being that current demographic literature predicts a negative impact of economic uncertainty on fertility rates (Sobotka et al. 2011; Vignoli et al. 2020). (H2) At higher inflation variance, the sensitivity of *λ*_*s*_ to the effect of inflation on survival is also higher. We hypotheised (H2) based on the findings that there exists a negative relationship between mortality and high inflation (Movsisyan et al. 2024) and the positive correlation between high inflation and inflation uncertainty (Fischer et al. 2024). Finally, (H3) high variability and autocorrelation in inflation rates adversely affect the capacity of human populations to buffer. We expect (H3) because the stochastic growth rate is negatively affected by temporal variances in vital rates and environmental autocorrelation following Tuljapurkar’s small noise approximation (Tuljapurkar 1982a, 1982b).

## Methods

We developed a quantitative framework using Bayesian Matrix Population Models (MPMs) to test our three hypotheses. Specifically, our framework comprised the following five steps: (1) extracting high-resolution inflation and vital rates data, (2) developing hierarchical Bayesian models for vital rates, (3) constructing MPMs using the posteriors from the Bayesian models, (4) computing stochastic growth rates, and (5) perturbation analysis of MPMs to examine the population response to variability in inflation environment. We performed all analyses in R v4.5.3 (R Core Team 2025), and our complete scripts are available at https://github.com/rmdemography/Inflation.

### Data

To test our hypotheses, we used inflation and demographic data from 32 countries spanning the period from 1971 to 2024. Specifically, we extracted “trend inflation” rates from *A Global Database of Inflation* (Ha et al. 2023), which contains global inflation estimates since 1970 for up to 209 countries. However, we used the annual “trend inflation” rates for 72 countries because the “trend” value is a long-term estimate of inflation that excludes transient variations (Stock and Watson 2016). Furthermore, we selected 32 countries for our analyses to ensure that each level of inflation and all geographic regions of the world were represented. A detailed table of the select countries, along with their mean and variance of trend inflation rates, is provided in the appendix.

We extracted the demographic data from the World Population Prospects (WPP) 2024 revision (United Naitons, Department of Economic and Social Affairs, Population Division 2024). The 2024 revision of WPP provides estimates and projections of key demographic indicators from 1950 to 2024 for 237 countries/areas. We used life table survival ratios and fertility rates to construct the Bayesian MPMs for the chosen 32 countries over the period from 1971 to 2024. Specifically, we used demographic data for females at single-year ages and at an annual frequency. Thus, we constructed female-only projection models. Moreover, we used the survival ratios from age 0 to 85 to avoid data quality issues at extreme ages, particularly in Low- and Middle-Income countries with poor data quality (United Nations, Department of Economic and Social Affairs, Population Division 2024).

### Matrix Population Models

To express the impact of inflation on population dynamics, we constructed inflation-dependent matrix population models. Specifically, we expressed the population structure as

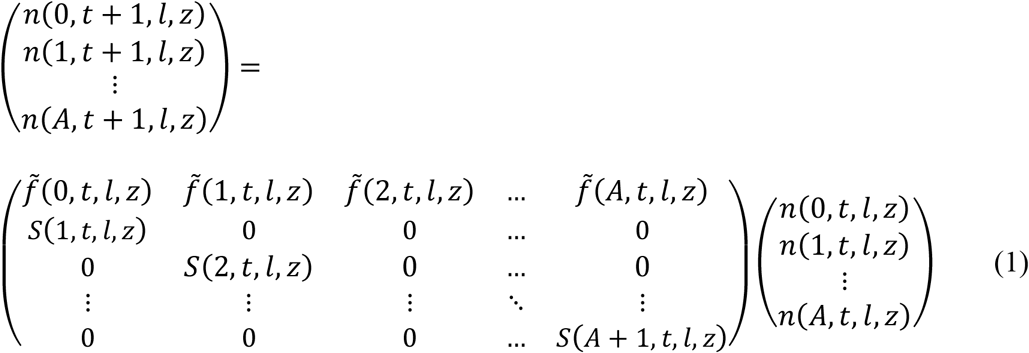

where, *n*(*x, t* + 1, *l, z*) represents the population aged *a* in country *l* at time *t* + 1, and the inflation level *z*. It is worth noting that we write *z* in parentheses in (1) because the vital rates and the MPMs are inflation-dependent, but that we drop the index *z* in the following sections for notational simplicity. The population grows from one age to the next in time *t* to *t* + 1 depending on the survival *S*(*x*) and fertility 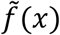 of population *n*(*x, t, l*) at age *x* and time *t*. To make the fertility rates sex specific and account for infant and maternal survival, we transformed the fertility rates as

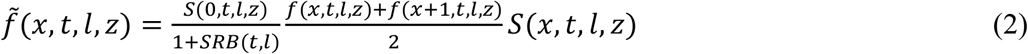

(Preston et al. 2001). Here, *f*(*x, t, l, z*) is the age-specific fertility rate, *SRB*(*t, l*) is the sex ratio at birth, *S*(0, *t, l, z*) is the infant survival ratio, and *S*(*x, t, l, z*) is the survival ratio at time *t*, country *l*, and inflation level *z*. Therefore, our MPMs are inflation-dependent, spatially and temporally stochastic.

### Model Specifications

To compute vital rates as a function of inflation rates, we estimate their posterior distributions using hierarchical Bayesian models. Specifically, we develop separate models for survival ratios *S*(*x, t, l, z*) and fertility rates *f*(*x, t, l, z*). Then we compute the posterior of 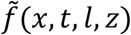 using 2.

#### Survival Model

For the survival model, we defined the logit-transformed survival ratios at age *a*, time *t*, and country *l* as

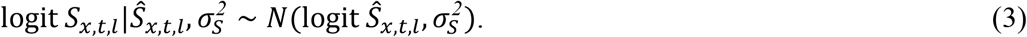

Here, the latent survival ratio is modelled on a logit scale as

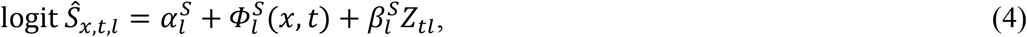

where 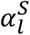 is the country-specific intercept, 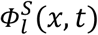 is the country-specific smoothened surface of survival over age and time, and 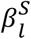 is the country-specific slope on log-transformed inflation rates *Z*_*t,l*_.

To estimate the baseline survival over age and time, keeping the marginal effect of inflation constant, we modelled the parameter 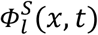 as a two-dimensional P-spline model. Two-dimensional P-spline models have been used in demographic literature to forecast mortality, with a second-order difference penalty to produce a smoothed age-time surface of mortality (Camarda 2019; Currie et al. 2004). Here, we assumed that the latent age-time surface of survival can be approximated by the tensor product of the two, one-dimensional B-splines,

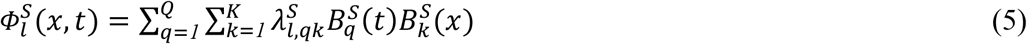

(Lang and Brezger 2004). We computed the cubic B-spline basis functions 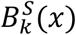 over age *x* and 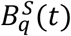 over time *t*, with eight equally spaced knots. Finally, to encode the smoothness of the two-dimensional survival surface over age and time, we model the country-specific basis coefficient 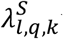 with an intrinsic conditional autoregressive (ICAR, hereafter) prior (Besag et al. 1991a; Besag and Kooperberg 1995).

To model the spline coefficient as an ICAR process, we followed the Besag-York-Mollie (Besag et al. 1991b) specifications. Specifically, we defined an unidirected graph on the two-dimensional *QK* lattice, with *Q* positions along the time-basis axis and *K* positions along the age-basis axis, by declaring two nodes adjacent if their coefficients are immediate neighbours along either axis. Hence, the set of edges is

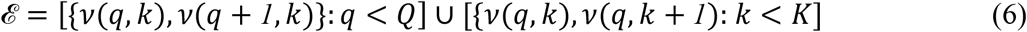

where *ν*(*q, k*) = (*k* − *1*)*Q* + *q*, is the position index on the vectorised lattice. Thereafter, we defined the joint distribution using the pairwise difference formulation

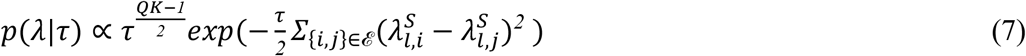

where *τ* is the precision parameter (M. Morris et al. 2019). To ensure the identifiability of the spline coefficient 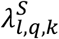, we added a sum-zero constraint 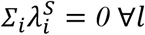. It is worth noting that our model is analogous to the Lang and Brezger (2005) Bayesian P-spline model, where the prior on spline coefficients is formulated as in Besag and Kooperberg (1995).

To make survival ratios inflation-dependent, we used log-transformed inflation rates *Z*_*t*−*2,l*_ at time *t* − *2* in country *l* as a covariate in the survival model. We used a two-year lagged inflation rate to account for the time-lagged effect of inflation on survival outcomes. Because previous studies exploring the impact of inflation on survival are scarce, we tested a two- to five-year lagged effect and ultimately chose a two-year lag. Additionally, there are multiple country-years with negative inflation rates (*i.e*., deflation; *e.g*., Argentina in 1998-2000 or India in 2019 and 2021[Ha et al. 2023]), and the distribution of inflation rates, especially in a few Latin American countries, is highly volatile with a large standard deviation (see Table A1 in the appendix). Therefore, we transformed the inflation rates to the continuously compounded scale using *log*(*1* + *r*_*t*−*1,l*_/*100*) where *r*_*t*−*2,l*_ is the two-year lagged inflation rate. We have also centred the transformed inflation rates within the country to ensure that the country-specific intercept 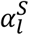 carries the intercept, 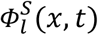 shapes the smooth age-time surface, and inflation rates provide additional deviation in the survival rates. Thereafter, we defined the country-specific slope 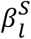 on inflation rates *Z*_*t,l*_ to account for the spatial heterogeneity in the inflation rates’ effect on survival.

#### Fertility Model

To estimate the posterior distribution of fertility rates as a function of inflation, we developed a hierarchical Bayesian model. Specifically, we defined the distribution of age-specific fertility rates *f*_*x,t,l*_ at time *t* in country *l* on a logit scale as

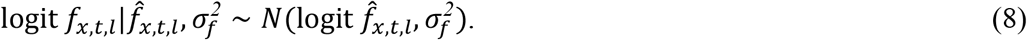

Then, we model the latent fertility rates as

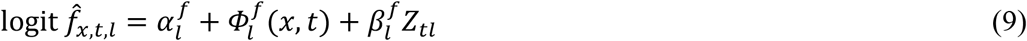

where 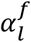 is the country-specific intercept, 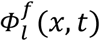 is the country-specific smoothened surface of fertility capturing how the age profile of fertility evolves jointly over time, and 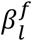 is the country-specific slope of log-transformed inflation rates *Z*_*t,l*_ on fertility rate. We incorporated log-transformed inflation rates *Z*_*t*−*2,l*_ as a covariate in the fertility model, as in the survival model. Here, we used two-year lagged inflation rates for the fertility model. Because, in line with previous literature (Adsera and Menendez 2011), the two-year lagged inflation rates on fertility give a better effect on fertility rates than three to five-year lagged inflation rates.

To encode a smooth age-time surface of fertility, we modelled the parameter 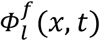 as a tensor product of penalised B-splines. Similar to equation (5) in the survival model, we constructed

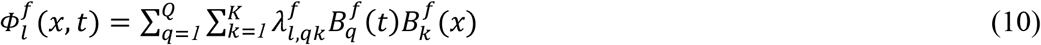

using a cubic B-spline basis 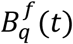 of dimension *Q* with eight equally spaced knots for the temporal dimension, and an analogous basis 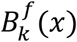 of dimension *K* over the age grid. We parameterised the smooth fertility surface for each country *l* by *QK* = *64* latent coefficients 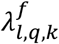, modelled as an ICAR process to penalise large differences in adjacent coefficients across the age-time grid (Besag and Kooperberg 1995; M. Morris et al. 2019). We defined the joint distribution for 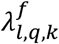 using (7), and the details of the priors are discussed in the subsequent section.

### Prior Specification

To complete the hierarchical Bayesian models for fertility and survival, we defined priors for the model parameters and hyperparameters. Specifically, we defined a non-informative prior on the country-specific intercept of fertility and survival as 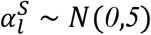 and 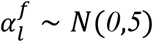, respectively.

For the precision hyperparameter *τ* in both fertility and survival models, we defined the prior

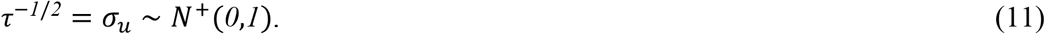

Morris et al. (2019) recommend a ∼ *G*(1,1) prior on *τ* for the stan implementation of the ICAR prior. However, the gamma prior for *τ* was creating a convergence problem for our model; therefore, we defined the half-normal prior on *σ*_*u*_. Additionally, we defined a hierarchical prior for the slope coefficient of inflation

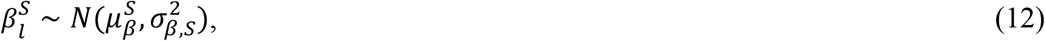

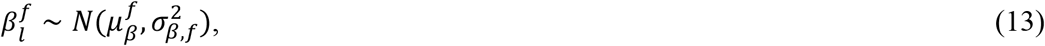

with 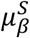 and 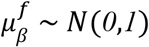. We completed the prior specification by placing a half-normal prior ∼ *N*^+^(*0,1*) on all the remaining *σ* parameters.

### Parameter Estimation

To obtain posterior distributions for the demographic parameters, we fit hierarchical Bayesian models using the Hamilton Monte Carlo (HMC) sampling algorithm. Specifically, for the computation of the proposed models, we used the HMC variant, the no-U-turn sampler (NUTS), which adapts algorithm parameters in each iteration to efficiently explore the posterior space and prevent inefficient trajectory reversals. The sampler is implemented using the “*cmdstanr”* interface (Gabry et al. 2025), which provides a robust HMC/NUTS-based implementation for Bayesian inference. For each model, we obtained four separate chains, each with 4000 iterations, out of which 2000 iterations were used as warm-up. We retained all iterations (without chain thinning) to estimate the associated posterior distribution.

We assessed model convergence using standard Stan diagnostic measures. First, we use the measure of split-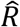 which compares the total variance to within-chain variance, representing the variability of the posterior samples within each chain. If the chains have not mixed well, the 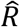 values become larger than 1. The values of 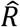 near 1 suggest that the between- and within-chain estimates are similar, *i.e*., all chains are exploring the same distribution. Therefore, Vehtari et al (Vehtari et al. 2021) recommend using the posterior samples only if the 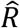 is less than 1.01. Second, we used the Effective Sample Size (ESS), which measures autocorrelation in the posterior draws, because HMC samples depend on the values from the last iteration. The ESS is calculated using the amount of autocorrelation, thereby allowing it to be used to diagnose sampling efficiency. We used two measures: Bulk ESS, estimated from rank-normalised draws, and Tail ESS, estimated as the minimum ESS at the 5% and 95% quantiles. Vehtari et al. (2021) suggest that both the bulk and tail-ESS should be at least 100 per chain to ensure sample reliability.

To construct the posterior distributions for inflation-dependent MPMs, we estimated the posterior distribution of survival (*S*_*a,t,l*_) and fertility (*f*_*a,t,l*_). Specifically, the hierarchical Bayesian models in (4) and (9) were used to obtain the posterior distributions. In turn, these models provided posteriors for the slope of inflation rates *β*_*l*_ per country, along with parameters *α*_*l*_ and *Φ*_*l*_(*x, t*) accounting for country-specific levels and the age-time surface of the vital rates. Therefore, we constructed the posterior distribution of MPMs from these parameters, making explicit the dependence of the MPM on inflation, along with uncertainty over country and time. Additionally, we performed the diagnostic tests to assess the reliability of the posterior samples, which showed that the models fit the data well. The split-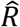 values for each parameter in the fertility and survival model were less than 1.01, indicating that the between- and within-chain variances are similar, and that both chains are exploring the same distribution. Furthermore, we found that both the bulk and tail effective sample sizes were higher than 400 (*i.e*., more than 100 times the number of chains), suggesting that posterior samples are reliable.

### Model Checks and Evaluation

To examine the validity of the Bayesian models for fertility and survival, we employed posterior predictive checks. Specifically, we simulate multiple replicates of fertility and survival rates using the samples drawn from the posterior predictive distribution and compare them with the observed rates. If the model is a good fit, it should generate replicates of rates that closely resemble the observed rates, and no systematic difference between the replicated and observed rates. We replicated the survival and fertility rates using the posterior predictive distribution as

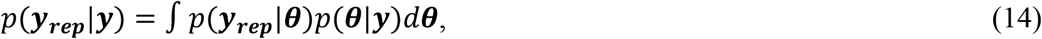

where ***y***_***rep***_ is the replicated rate, ***y*** is the observed rate, and ***θ*** contains the model parameters. We assessed model fit by overlaying the observed rate density with the simulated rate density (see Appendix).

Additionally, we compared the replicated rates with the observed rates using test statistics *T*[***y, θ***], which can be defined as a scalar summary of the rates. With no clear guidance on selecting the test statistics, we computed the mean and standard deviation of the rates. Thereafter, we constructed histograms based on the mean and standard deviation for each replicated dataset. For a good-fit model, the test statistic for the observed rate should lie at the middle of the histogram; if the test statistics lie at the outer ends of the histogram, it indicates that the model cannot reproduce the observed rates. The figure for each test statistic for both the fertility and survival models is given in the appendix.

Finally, we evaluated the model using mean absolute error (MAE) and root mean squared error (RMSE). The details of the MAE and RMSE for both the survival and fertility models are given in the appendix.

### Simulate the Inflation Environment

To explore the roles of variability in inflation on population dynamics, we simulated the MPMs across the environmental autocorrelation-variance parameter space. Specifically, we estimated a 1000-year time-series of inflation rates for all combinations of stochastic environmental parameters (*i.e*., *φ* and *σ*), with autocorrelation *φ* ranging from 0.5 to 0.9 and standard deviation *σ* ranging from 0.1 to 0.5. We used a first-order autoregressive (AR(1), hereafter) function to generate the sequence of inflation rates used to build the series of MPM elements

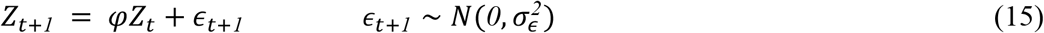

where *φ* represents the degree of autocorrelation across time steps and *ϵ*_*t*+*1*_ represents the white noise. We incorporated the standard deviation of inflation rates into the AR(1) simulation as

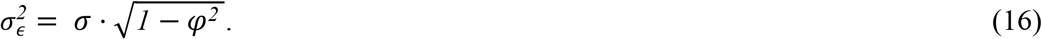

Thereafter, we used the simulated series of inflation rates, along with other posterior draws, to construct the stylised MPMs for the 1,000 {Citation}time steps. Generating these inflation time series subsequently creates a parameter space of (inflation) environmental stochasticity with axes of temporal autocorrelation and variance. It is worth noting that this parameter space does not represent a realised scenario in nature, as environmental variables do not vary with the specified variance and autocorrelation for our population of interest. The purpose of the parameter space is to manipulate the degrees of environmental stochasticity across two axes to inform how vital rates respond to inflation uncertainty.

### Reparameterise the MPMs

To construct the inflation-dependent Bayesian MPM, we parameterised (1) using posteriors from the Bayesian models. Specifically, we extracted the posterior draws of the survival ratio 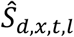 and fertility rates 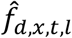, and then computed the posteriors of 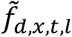 using equation 2. Following standard demographic conventions, we set the SRB to 1.05 (Wheldon et al. 2013). Note that the index *d* in subscript indicates the specific posterior draws. Finally, using the posterior distributions of survival 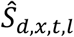 and fertility 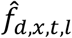, we construct the Leslie matrix, 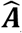, for the projection model given in (1). In summary, we obtain posteriors for the projected population structure, with the draws representing the uncertainty in the estimates.

We built posteriors for the MPMs over each simulated inflation time series using five steps. First, we drew posteriors of time-invariant intercept *α*_*l*_ and slope coefficient *β*_*l*_ from both the survival (4) and the fertility model (9). Second, we constructed posteriors of *Φ*_*l*_(*x, t*) using the spline bases *B*_*k*_(*x*) and *B*_*q*_(*t*), and the coefficient *λ*_*l,q,k*_. Note that the spline coefficients are defined for the age and time basis evaluated over the examined age and year range; hence, the coefficients would not be valid if we recomputed the time basis *B*_*q*_(*t*) over the simulated time range. Therefore, we fixed the time basis at the last knot *B*_*Q*_(*t*), assuming that the age-time surface of the vital rates remains similar to the later part of observed years over the simulated time series and temporal variance in the vital rates would be produced by the variability in inflation rates. Third, we collected the simulated series of inflation rates over 1000 timesteps for each combination of *φ* − *σ* values. Fourth, we estimate the posteriors for survival and fertility rates over the simulated series using

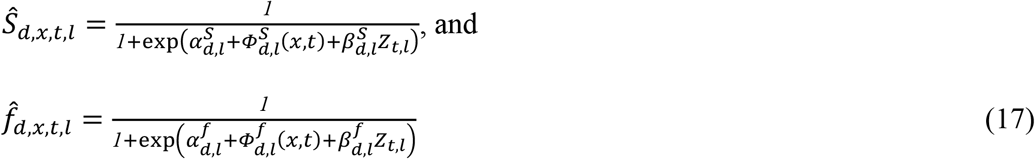

Fifth, we computed the 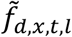 using (2) and computed posteriors of MPMs *Â*(*d, t, l*) over the 1000 simulated time steps for each combination of *φ* − *σ* by using 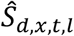 and 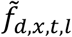 in (1).

### Stochastic Growth Rate

To estimate the long-term stochastic population growth *λ*_*s*_ dependent on inflation rates, we derived the growth rate *λ*_*s*_ from the posterior distribution of vital rate parameters. The stochastic population growth rate is given by the long-run geometric mean of annual growth rates

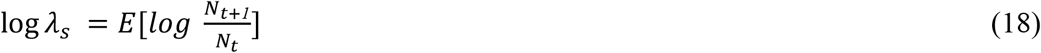

where *N*_*t*_ is the population size, and *E* gives the expected value. Whilst methods exist to analytically compute the stochastic growth rate *λ*_*s*_ (Tuljapurkar 1982a, 1982b), we used a numerical simulation in our analysis. Specifically, to calculate the posterior distribution of the *λ*_*s*_, we extracted the posteriors for the MPMs from the previous step. Thereafter, for each draw, we simulated the population dynamics for 1000 years and calculated the geometric mean of the *N*_*t*+*1*_ = *Â*_*t*_*N*_*t*_ (analogous to the approach in Elderd and Miller 2016). Hence, for a single parameter set, MPM elements 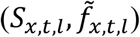 varied over time based on the variability of the simulated time series of inflation rates and across countries based on random draws from the country-specific model parameters. Therefore, the distribution of *λ*_*s*_ across parameter sets reflects the total uncertainty in the population growth rate, including estimation error of the vital rate coefficients and two types of process errors: country and time-specific variances in vital rates.

### Sensitivity Analysis

To analyse the effect of inflation rate variability on the growth rate, we performed sensitivity analyses of the growth rate to perturbations in the inflation slope coefficient. We examined the effect of perturbing the country-specific slope coefficient *β*_*l*_, which describes the inflation-dependent survival ratios and fertility rates. For each combination of environmental parameters (*φ, σ*), country, and posterior draw, we compute the sensitivity of the stochastic growth rate to a perturbation in the slope coefficient *β*_*l*_ as

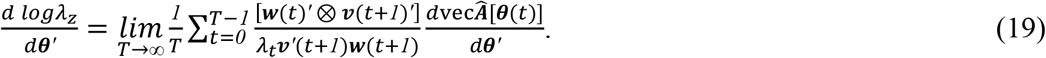

Here, ***w***(*t*) is the stable age distribution, ***v***(*t*) is the reproductive value vector, and *λ*_*t*_ is the growth of total population size from *t* to *t* + *1*. Note that ***θ*** contains the parameters from the Bayesian hierarchical models in (4) and (9), and we obtain the sensitivity by computing the derivative of 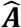 with respect to the perturbation in the slope coefficient *β*_*l*_. For notational simplicity, we did not mention all indices for elements in the sensitivity formulation in (19). Moreover, to test our hypothesis, we estimated the stochastic sensitivity of the stochastic growth rate *λ*_*s*_ to perturbations in the slope coefficient of inflation on fertility 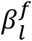 and survival 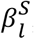, respectively. Interestingly, the elasticities with respect to *β*_*l*_ are less clearly interpretable than the corresponding sensitivities, as the slope coefficient is negative and a proportional change would make it even more negative. Whereas, stochastic sensitivities play an important role in evolutionary demography as they have the opposite sign to elasticity and can be interpreted as a fitness gradient for evolutionary stability analysis (Rees and Ellner 2009). Therefore, for negative underlying parameters, sensitivity rather than elasticity will correctly indicate the predicted direction of long-term adaptive dynamics.

### Elasticity Analysis

To quantify the influence of inflation uncertainty on the degree of demographic buffering, we calculated the sum of stochastic elasticities of variance in vital rates with respect to *λ*_*s*_ across the *φ* − *σ* space. We estimate this variable, 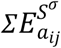, numerically. Since our matrix *Â* for MPM in (1) is composed of individual matrix elements *a*_*i,j*_ of survival and fertility, and is simulated for 1000 time-steps across the temporal autocorrelation (*φ*)-standard deviation (*σ*) space, we can quantify the temporal variance of each matrix element *a*_*ij*_ in the series of 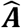 matrices in the space. The sum of stochastic elasticity quantifies the extent to which the stochastic growth rate is influenced by changes in the variances of the MPM elements (*i.e*., fertility and survival). A lower absolute value of the sum indicates that the stochastic growth rate is less sensitive to variance in the MPM elements, *i.e*., that the variance is constrained by natural selection, supporting DBH (Haridas and Tuljapurkar 2005).

To compute the sum of stochastic elasticity with respect to variance 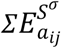, we followed Tuljapurkar et al. (Tuljapurkar et al. 2003). Specifically, we computed the derivatives

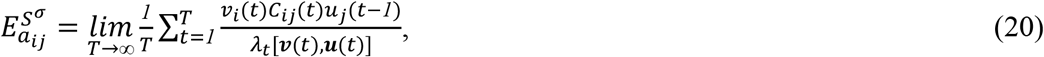

where the perturbation 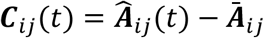, increases the variance of the (*i, j*) rate by a factor (*1* + *δ*)^*2*^ without changing the means. The number *δ* is the proportional increase in the standard deviation *σ*_*ij*_ of the vital rates and *Ā* is the average matrix over the simulated time series. Hence, we obtained the elasticity with respect to standard deviation (variance) as

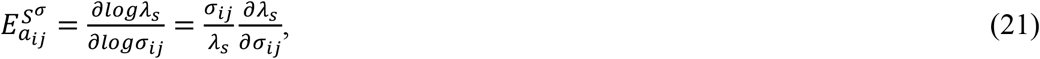

and computed the sum 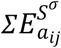 to quantify demographic buffering (Santos et al. 2024; Tuljapurkar et al. 2003). Furthermore, we performed the analyses for each of the MPM posterior distributions constructed above. Therefore, the distributions of the sum of stochastic elasticities with respect to variance across the *φ* − *σ* parameter space indicate the effect of inflation uncertainty on the degree of demographic buffering, and their distributions across the posterior sets represent the uncertainty in the estimates due to spatio-temporal fluctuations in vital rates.

## Results

We find support for our hypothesis H1, that the sensitivity of the growth rate *λ*_*s*_ to the slope coefficient of inflation on fertility 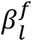 increases with inflation variance and autocorrelation. Specifically, we estimated the stochastic sensitivity 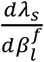 at each point in the standard deviation *σ*-autocorrelation *φ* parameter space of the inflation environment. In support of H1, we find that as standard deviation and temporal autocorrelation in log-transformed inflation rates increase, stochastic sensitivity increases across all examined countries (see Figure A9 in the appendix). Additionally, interesting patterns are observed on close inspection of the sign of the slope and the sensitivity across the *σ* − *φ* parameter grid. In countries including Chile, Colombia, Indonesia, Malaysia, Peru, Singapore, Tunisia, and Venezuela, the stochastic sensitivity is higher and negative at higher *σ, φ* values, indicating that increasing the negative slope coefficient, *i.e*., taking it closer to zero, reduces the stochastic growth rate (see Figure 1). On the other hand, in Argentina, Ecuador, the Philippines, Sweden, and Thailand, the sensitivity of the stochastic growth rate to a negative slope coefficient is positive and higher at high values of environmental parameters (*σ, φ*). The result indicates that taking the negative slope coefficient 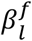 closer to zero increases the stochastic growth rate. Interestingly, in three out of the 32 examined countries, including Finland, Norway, and Taiwan, the slope coefficient is positive and shows a high and positive sensitivity at higher values of the *σ* − *φ* grid. However, it is worth noting that the slope coefficient of inflation on fertility across all the examined countries is small in magnitude, and the sensitivity of the stochastic growth rate 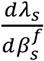 is also closer to zero, showing no uniform pattern in the sign of the sensitivity.

**Figure 1.**
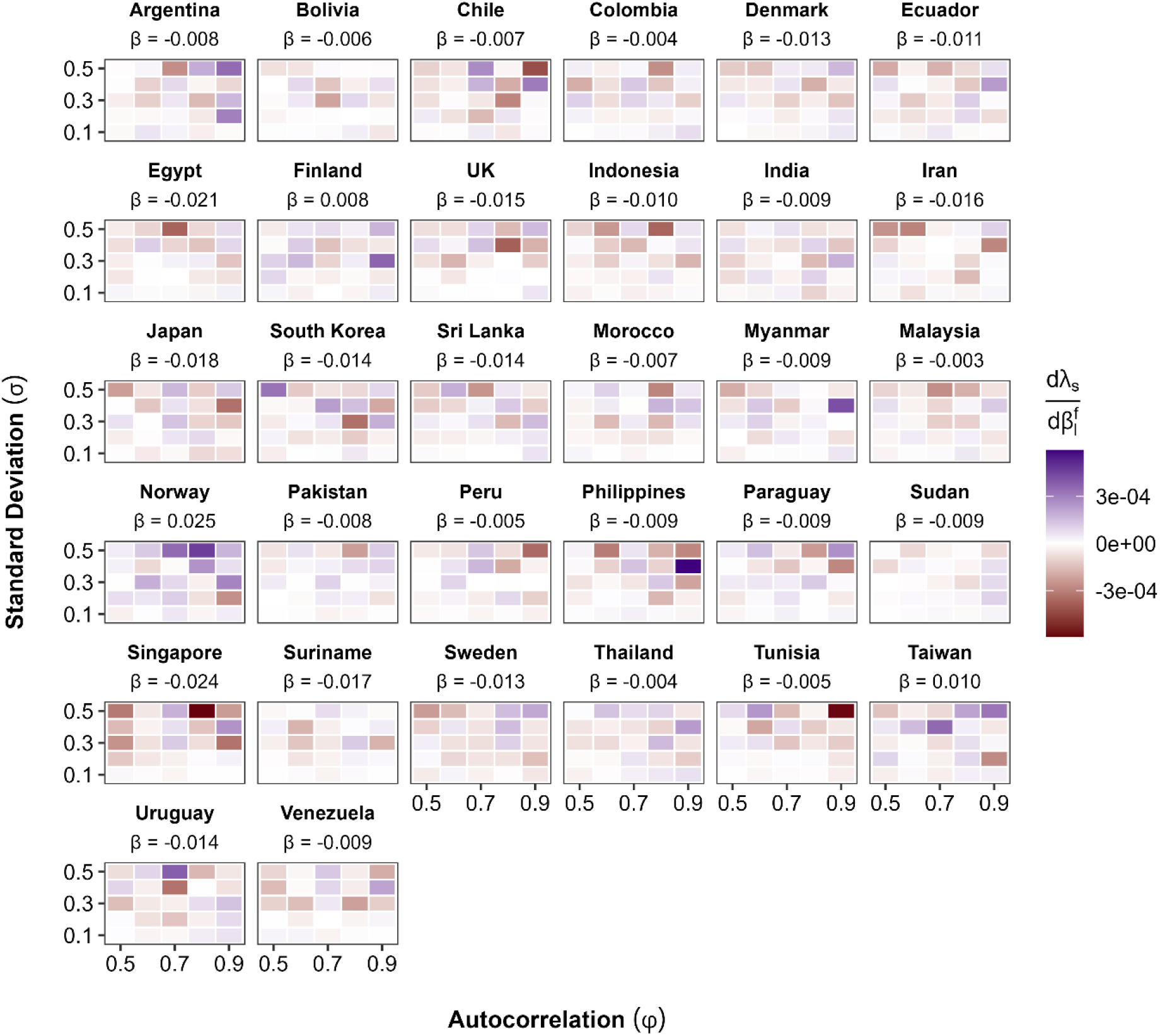
The sensitivity of stochastic growth rate to the slope coefficient of inflation on fertility 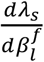 increases with variance and autocorrelation in inflation. The figure presents the mean of the posterior distribution of the stochastic sensitivity 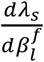. Positive sensitivities are shown in reds and the negative sensitivities are shown in purple across the *σ* − *ρ* plane. No unique pattern in the sign of sensitivities is observed, and the pattern of increase in 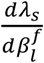 with increasing *σ* and *ρ* varies across countries.

Similarly, we found support for the hypothesis (H2) that with increasing uncertainty in inflation rates, fluctuations in survival can translate into fluctuations in the stochastic growth rate *λ*_*s*_. We find that the sensitivity of the stochastic growth rate to the slope coefficient of inflation rates on survival ratios 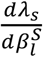 increases with increasing values of environmental parameters (*σ, φ*) across all the examined countries (see Figure A10 in the appendix). In countries including Colombia, Egypt, Finland, Srilanka, Norway, and Uruguay, at higher values of the *σ* − *φ* grid, the stochastic sensitivity is large and negative, indicating the fact that increasing the negative slope coefficient 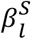, *i.e*., taking it closer to zero, reduces the stochastic population growth rate (see Figure 2). On the other hand, in Bolivia, Ecuador, Indonesia, Japan, Malaysia, Pakistan, Peru, Philippines, Paraguay, and Tunisia, the stochastic sensitivity to the negative slope coefficient 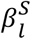 is positive at higher values of *σ* and *φ* reflecting the opposite scenario that taking the slope coefficient closer to zero increases the stochastic growth rate. Interestingly, in Argentina, South Korea, Myanmar, Sudan, Suriname, Sweden, and Venezuela, the slope coefficient 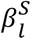 is positive and the stochastic sensitivity at higher values of *σ* − *φ* the grid is also high but negative. Compared with fertility, both the slope coefficient and the stochastic sensitivity are much smaller in magnitude. Moreover, most of the sensitivity values are very close to zero, and similar to fertility, do not show any unique pattern in the sign of the sensitivity values with increasing *σ* and *φ*.

**Figure 2.**
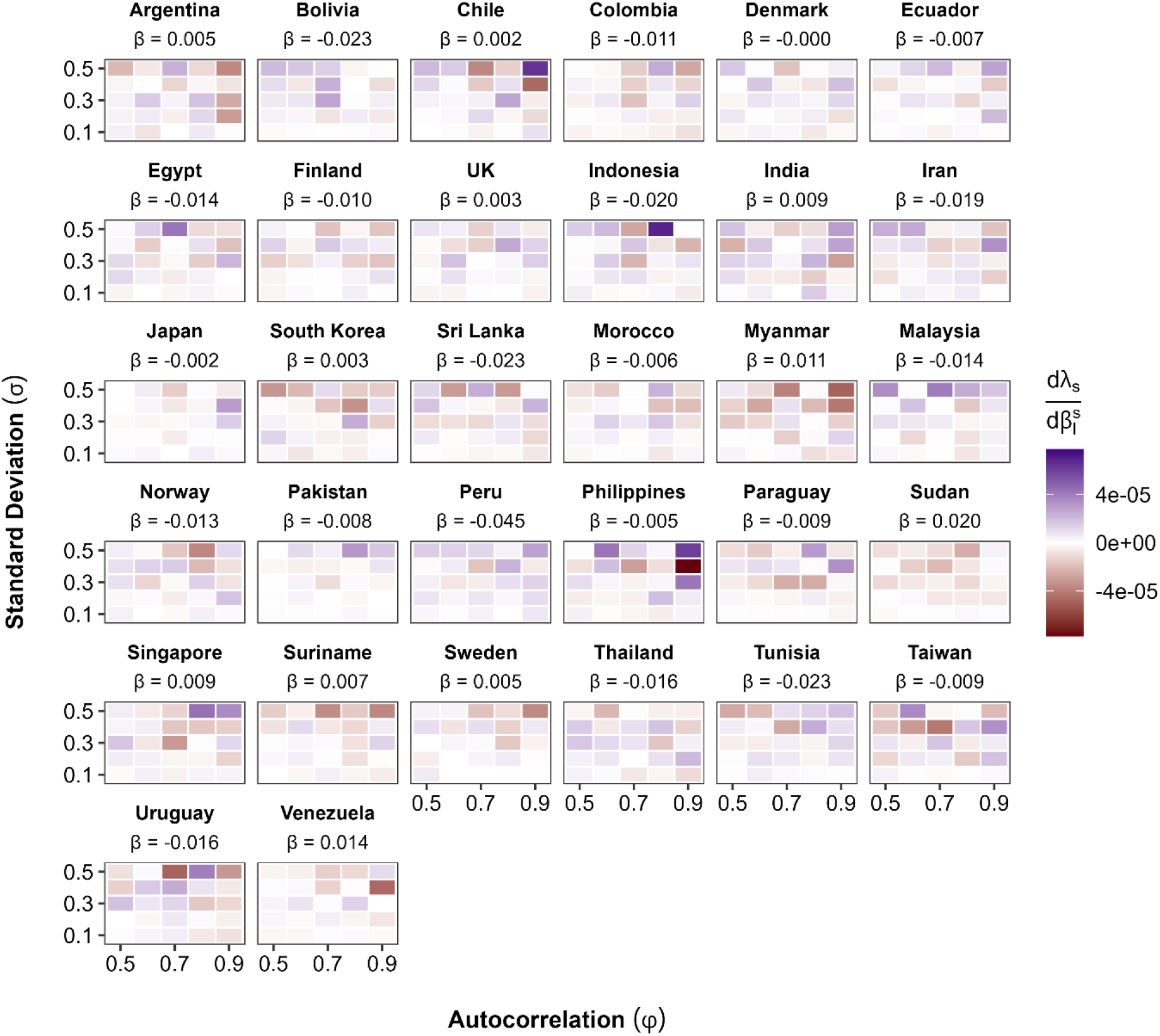
The sensitivity of stochastic growth rate to slope coefficient of inflation on survival 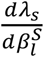 increases with variance and autocorrelation in inflation rates. The figure shows the mean of the posterior distribution of the stochastic sensitivity 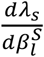. Positive sensitivities are shown in reds and the negative sensitivities are shown in purple across the *σ* − *ρ* plane. Furthermore, 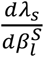 is much lower than the sensitivity for fertility 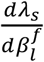 for all examined countries.

We do not find support for our hypothesis (H3) that with increasing inflation variance and temporal autocorrelation, the capacity to buffer decreases among human populations. Interestingly, we find that in most of the examined countries, including Bolivia, Colombia, Finland, Indonesia, Finland, Morocco, Malaysia, Pakistan, Peru, Paraguay, Sudan, Thailand, Tunisia, Uruguay, and Venezuela, the sum of stochastic elasticity with respect to variance in vital rate 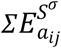 is approximately zero across the *σ* − *φ* parameter space (See Figure 3). Although in Argentina, Denmark, Ecuador, Myanmar, and Sweden, we find a smaller 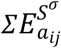 at a higher grid value, the values are very small to show any visible pattern supporting our hypothesis (H3).

**Figure 3.**
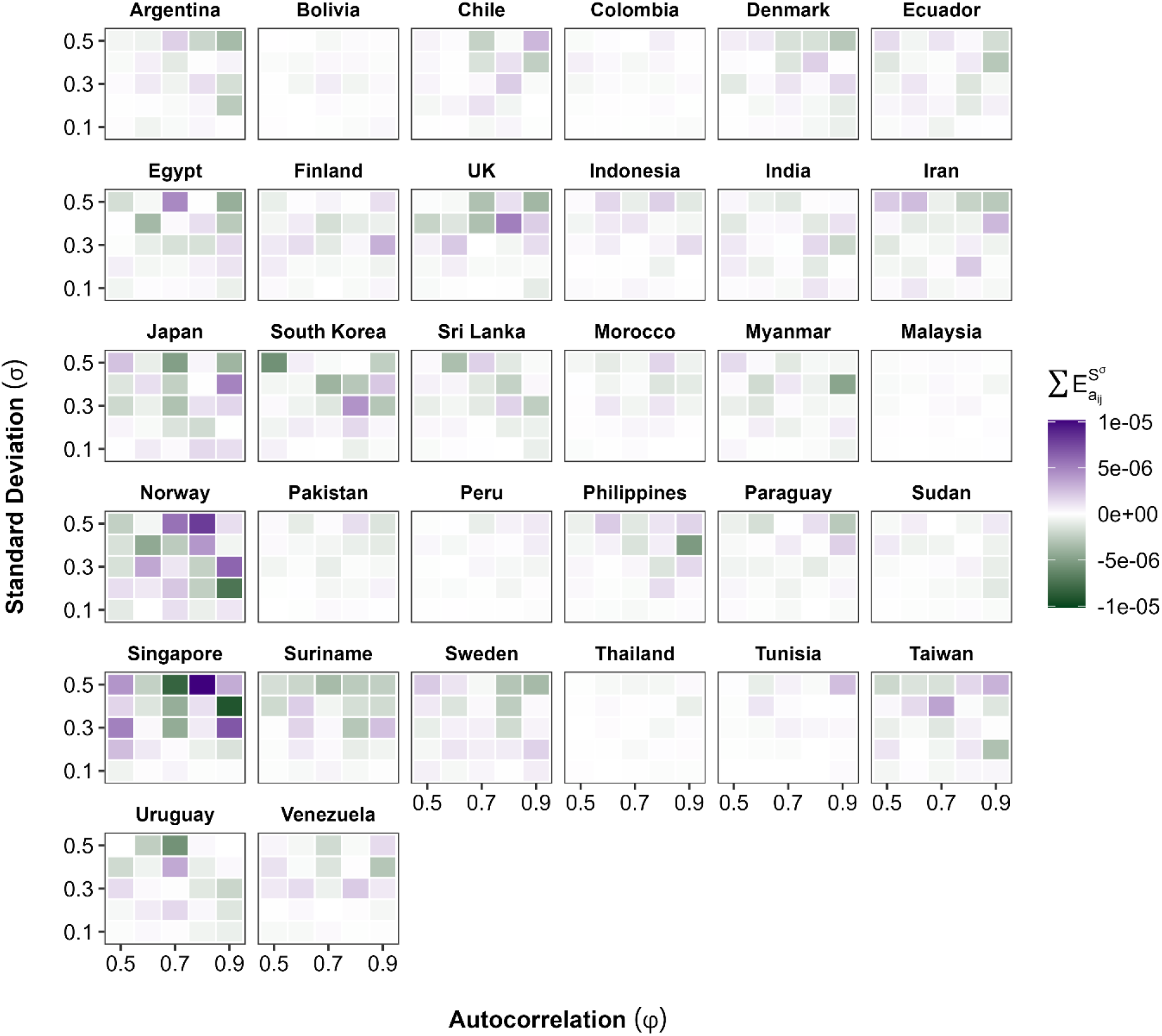
The capacity for demographic buffering remains approximately constant increasing variance and autocorrelation in most of the examined countries. Sum of stochastic elasticity of long-term growth rate *λ* to variance in vital rates 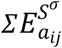 used as the measure of demographic buffering. Here, we present the mean of 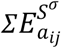 distribution computed from the posterior sample of vital rates and *λ*_*s*_.

## Discussion

Recurring evidence suggests that economic uncertainty shapes key demographic decisions, making the examination of inflation uncertainty increasingly important. In particular, inflation uncertainty can influence investment in fertility and the trade-off between survival and fertility, raising the question of how inflation variance affects humans’ capacity to buffer against a variable economic environment. To address this knowledge gap, we used hierarchical Bayesian models to parameterise long-term human survival and fertility data across 32 countries, enabling us to examine how inflation uncertainty shapes human population dynamics. Specifically, our models implemented with the log-transformed inflation rate as a covariate, accounting for the uncertainty associated with (spatiotemporal) process and measurement error, show the slope of inflation on each vital rate and propagate the uncertainty into a probability distribution for the MPMs and quantities derived from them (analogous to the results for IPMs in Elderd and Miller 2016; Rees and Ellner 2009). As humans are a relatively long-lived species (Jones et al. 2014) with complex life histories that respond to spatio-temporal variance in the economy, technological advancements, climate, and stochastic events such as pandemics, war, and natural disasters (Hellstrand et al. 2024; Vignoli et al. 2020; Vollset et al. 2024), explicit consideration of spatio-temporal variance in vital rates is essential for understanding the impact of inflation. Using Bayesian MPMs, we find support for (H1), suggesting a perturbation in the marginal effect of inflation on fertility can lead to a fluctuation in the stochastic growth rate. Similarly, we find an increase in the stochastic sensitivity to the slope coefficient on survival with increasing variance and autocorrelation in the inflation environment, but its smaller magnitude suggests that humans attempt to buffer survival against variable inflation, potentially by trading off fertility. However, we do not find support for the hypothesis (H3) that humans’ capacity to buffer is adversely affected by inflation variance and autocorrelation.

Stochastic sensitivities of long-term population growth rate to perturbation in the slope coefficient 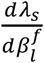 indicate directional selection on the slope coefficient. Our results show that, at higher inflation variance and autocorrelation, taking the negative slope of inflation on fertility closer to zero reduces the stochastic population growth rate in Chile, Colombia, Indonesia, Malaysia, Peru, Singapore, Tunisia, and Venezuela. The result indicates that the relative gain in fertility during low-inflation years contributes more to the stochastic growth rate than the fertility loss in high-inflation years in these countries, and the perturbation that weakens the negative inflation-fertility relationship reduces the stochastic growth rate. Our findings support the argument that the population can benefit by paying attention to the environment and reproducing effectively under each condition (Orzack 1985). Interestingly, we observed the opposite phenomenon in Argentina, Ecuador, the Philippines, Sweden, and Thailand, where the negative slope of inflation on fertility is under positive selection; *i.e*., taking the negative coefficient closer to zero would increase the stochastic growth rate. Here, the relative loss in fertility during the high-inflation years dominates the long-term stochastic growth rate, and hence, taking the negative slope closer to zero would increase the growth rate. However, it is worth noting that the sensitivity of the stochastic growth rate to perturbations in the slope coefficient of inflation on fertility 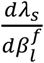 is very small in magnitude across the 32 countries examined. Hence, fluctuations in inflation, even at the highest level of the environmental parameter (*σ, φ*), do not lead to noticeable fluctuations in the stochastic population growth of humans.

Stochastic sensitivity to the inflation slope coefficient on survival 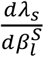 provides similar insights to those in fertility. Particularly in environments with high inflation variance and autocorrelation, perturbations in the negative slope of inflation can reduce the stochastic growth rate in countries such as Colombia, Egypt, Finland, Sri Lanka, Norway, and Uruguay. In these countries, the relative gain in survival in the low-inflation years over the long-term inflation trajectory outweighs the relative reduction in survival during the high-inflation years; therefore, weakening the negative relationship between inflation and survival would reduce the stochastic growth rate. Whereas in Bolivia, Ecuador, Indonesia, Japan, Malaysia, Pakistan, Peru, Philippines, Paraguay, and Tunisia, the relative reduction in survival in high-inflation years mostly determines the stochastic growth rate, and taking the negative slope coefficient 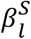 closer to zero would increase the stochastic growth rate. However, the stochastic sensitivity 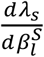 is very close to zero and much smaller in magnitude than the sensitivity for the case of fertility 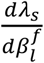 suggesting that the parameter 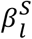 is close to an evolutionarily stable strategy, at which there is negligible or no directional selection (Rees and Ellner 2009). We speculate that the evolutionary stable strategy for the slope parameter on survival 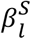 occurs because if there is a trade-off between investment in survival and fertility, humans would choose survival over fertility. Our speculation is based on the fact that, despite stochastic events such as war, epidemics, economic depressions, or groundbreaking medical inventions, age-specific survival among humans has been increasing at a remarkably constant pace (Lee and Carter 1992; Tuljapurkar et al. 2000; Wachter 2003).

The stochastic sensitivity to the slope of inflation rates, simulated across a variance-autocorrelation parameter space, quantifies the impact of inflation uncertainty. Variance in the inflation environment can affect the underlying vital rates of the Bayesian MPM, parameterised as a continuous function of the inflation rates, and thereby influence stochastic population growth (Gascoigne et al. 2025; Haridas and Tuljapurkar 2005). Additionally, environmental autocorrelation can have a significant impact on the stochastic population growth rate, which is jointly determined by environmental memory (i.e., the magnitude of autocorrelation) and demographic damping. As a result, populations with weaker damping tend to remember the variability in vital rates and accumulate it over time (Gascoigne et al. 2025; Tuljapurkar and Haridas 2006). In line with these arguments, our results suggest that as the variance and serial autocorrelation in the inflation environment increase, the sensitivity of stochastic growth rates to the slope parameters of fertility 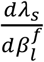 (H1) and survival 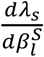 (H2) increases across all the examined countries. However, the magnitudes of the sensitivities for both fertility and survival are very close to zero, even at high values of the (inflation) environmental variance and autocorrelation. Additionally, the signs of the stochastic sensitivities do not show a uniform pattern across countries. Therefore, we infer that fluctuations in inflation rates across the examined countries do not lead to large fluctuations in the stochastic growth rates through changes in the underlying vital rates.

Examining the effects of inflation variance and autocorrelation on humans’ capacity for demographic buffering yields valuable evolutionary insights. The demographic buffering quantified by the sum of stochastic elasticity 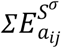 with respect to variance in vital rates in the Bayesian MPMs does not support the hypothesis (H3) that increasing variance and autocorrelation in the inflation environment negatively affect humans’ buffering capacity. Although existing ecological literature suggests that environmental variance negatively affects buffering through its impact on vital rates, while autocorrelation affects through a shift in population structure may have a curvilinear effect (Gascoigne et al. 2025). Our results show that, in almost all the examined countries, the capacity to buffer does not decrease with increasing variance and temporal autocorrelation in inflation levels. The result supports the fact that the slope of inflation on fertility and survival across the examined countries is very small in magnitude, so human vital rates might not respond strongly to fluctuations in national-level inflation rates. Additionally, we speculate that human populations, especially in countries with a historically high mean inflation rate, have developed a life history strategy against inflation uncertainty that does not adversely affect their capacity to buffer. In line with this argument, empirical evidence shows that in some of the abovementioned Latin American countries with very high mean inflation and frequent episodes of inflationary shocks, women utilise a temporary period of unemployment as a favourable time for conception and childbearing (Adsera and Menendez 2011).

Certain notable limitations of our study call for a cautious interpretation of the results. Specifically, due to the limitations of sex-specific migration data in the WPP, we do not consider migration as a vital rate in our inflation-dependent MPMs, and therefore in the stochastic population growth (United Naitons, Department of Economic and Social Affairs, Population Division 2024). As our objective is to examine the selection gradient on the slope of inflation on fertility and survival, the computed long-term stochastic growth indicates the population growth that we would observe if the population experienced the simulated vital rates of fertility and survival, along with zero net migration. However, inflation uncertainty significantly affects migration rates (Rocha et al. 2022; Salisu et al. 2024), and examining the selection gradient on migration rates is a promising avenue for future research. Additionally, we utilised the demographic estimates of survival and fertility from the WPP 2024 version; therefore, our results are sensitive to the assumptions of the WPP methodology (United Nations, Department of Economic and Social Affairs, Population Division 2024). It is also worth noting that our MPMs are female-dominant models. A two-sex extension of our models can be easily constructed by following the two-sex model for demographic reconstruction developed by Wheldon et al. (2015); however, it requires substantial computational resources. Finally, our models can be referred to as the “mean-field” description of large human populations, for which the effect of demographic stochasticity as an underlying mechanism of buffering can be ignored (Rees and Ellner 2009). Therefore, future studies should examine the models in which the mean field model is perturbed by random variables approximating the net effect of individual-level random events, *i.e*., demographic stochasticity.

Our framework provides a valuable tool for examining the evolutionary impact of environmental drivers on life history strategies within a stochastic environment. Specifically, the hierarchical Bayesian approach for constructing the stochastic MPMs models the demographic rates as continuous functions of the underlying (continuous) inflation rates and models the nonlinear spatiotemporal variations by incorporating continuous prior distributions for the parameters defining the demographic rates (Elderd and Miller 2016; Rees and Ellner 2009). The Bayesian MPMs quantify inflation-dependent demographic rates while accounting for spatio-temporal heterogeneity, covariance, and uncertainty (process error) in vital rates, which in turn translates into uncertainty in population-level inferences about stochastic population growth rates, sensitivity, and elasticity. Interestingly, the hierarchical Bayesian framework for MPMs offers the flexibility to utilise data from multiple sources, such as the different sources of inflation and demographic data in our study, while accounting for the various sources of measurement error associated with the data (Elderd and Miller 2016; Rees and Ellner 2009). Most importantly, the parametric elements of the Bayesian MPM smoothly fitted as a continuous function of underlying parameters, *i.e*., slope coefficient (of inflation rates) in the Bayesian models, enables us to perturb the parameter and analyse the stochastic sensitivity of the growth rates, which can be interpreted as the fitness gradient for evolutionary stability analysis (Rees and Ellner 2009). Therefore, our framework can be applied to any environmentally explicit structural population to estimate the robust probability distributions for stochastic population growth and its sensitivities and elasticities, which play a crucial role in evolutionary demography.

In conclusion, our study makes a key contribution to the existing literature examining the relationship between economic uncertainty and demographic behaviour. Building on the theory of stochastic demography (Rees and Ellner 2009; Tuljapurkar 1989), we perform a perturbation analysis of the potential response of long-term population dynamics to inflation uncertainty within stochastic environments. We present novel findings indicating that the slope of inflation on vital rates, including fertility and survival, is subject to weak selection, and the selection pressure increases with the variance and serial autocorrelation of inflation rates. In line with the classic conclusion from the studies on human longevity (Lee and Carter 1992; Tuljapurkar et al. 2000; Wachter 2003), suggesting that humans preserve the shape of survival even in the stochastic conditions, we find that the survival rates of human populations are near an evolutionary stable strategy when examined against inflation uncertainty within variable environments. Additionally, we find that humans’ capacity to buffer remains largely unaffected by increasing variance and autocorrelation in the inflation environment. Notwithstanding the novelty of the empirical evidence, our study develops a robust quantitative framework to examine the evolutionary impact of any environmental drivers on the life history strategies among humans within stochastic environments.

## Acknowledgement

We thank Jennifer B Dowd for her helpful feedback on this manuscript. RSG was supported by a NERC Pushing the Frontiers grant (NE/X013766/1).

