## Supplementary appendix for "Humans exhibit demographic buffering against variability and persistence in inflation environments"

### Variability and persistence in inflation reshape human population dynamics by amplifying sensitivity and eroding demographic buffering

Rahul Mondal[1,*], Samuel JL Gascoigne[2], Udaya S Mishra[1], Roberto Salguero-Gomez[3,*]

#### Appendix

**Table 1.** Descriptive statistics of the inflation in rates across the 32 examined countries during the period between 1971-2021.

| Sl. No. | Country | Mean | SD | Median | Min. | Max. |
|---|---|---|---|---|---|---|
| 1 | Argentina | 104.89 | 155.18 | 29.66 | -1.08 | 738.78 |
| 2 | Bolivia | 25.33 | 42.92 | 7.35 | 0.41 | 179.5 |
| 3 | Chile | 42.85 | 109.46 | 7.93 | -0.19 | 633.85 |
| 4 | Taiwan | 2.63 | 2.94 | 1.67 | -1.58 | 14.08 |
| 5 | Colombia | 14.86 | 9.5 | 16.22 | 1.94 | 31.56 |
| 6 | Denmark | 4.22 | 3.84 | 2.46 | -0.38 | 13.87 |
| 7 | Ecuador | 20.7 | 21.96 | 11.85 | -0.03 | 93.2 |
| 8 | Egypt | 10.54 | 5.76 | 10.27 | 2.38 | 23.91 |
| 9 | Finland | 4.49 | 4.6 | 3 | -0.11 | 18.26 |
| 10 | India | 7.3 | 5.47 | 7.85 | -10.03 | 26.08 |
| 11 | Indonesia | 9.27 | 7.08 | 8.3 | 0.97 | 45.51 |
| 12 | Iran | 18 | 9.13 | 17.83 | 1.91 | 42.74 |
| 13 | Japan | 2.08 | 3.55 | 0.54 | -0.71 | 16.84 |
| 14 | Malaysia | 3.36 | 3.14 | 2.88 | 0.03 | 21.05 |
| 15 | Morocco | 4.43 | 4.26 | 2.85 | -1.25 | 16.64 |
| 16 | Myanmar | 14.26 | 15.84 | 6.83 | -10.49 | 58.26 |

| Sl. No. | Country | Mean | SD | Median | Min. | Max. |
|---|---|---|---|---|---|---|
| 17 | Norway | 4.35 | 3.22 | 2.6 | 1.18 | 13.24 |
| 18 | Pakistan | 8.76 | 5.51 | 7.64 | 2.6 | 31.28 |
| 19 | Paraguay | 12 | 9.47 | 8.22 | 1.99 | 39.19 |
| 20 | Peru | 33.25 | 41.01 | 7.82 | -0.17 | 120.55 |
| 21 | Philippines | 9.35 | 8.59 | 7.32 | -0.99 | 40.44 |
| 22 | Republic of Korea | 3.86 | 2.85 | 3 | 0.51 | 14.36 |
| 23 | Singapore | 0.77 | 1.22 | 0.5 | -0.61 | 6.77 |
| 24 | Sri Lanka | 9.33 | 6.1 | 8.99 | -3.28 | 22.2 |
| 25 | Sudan | 39.01 | 39.47 | 23.66 | 1.11 | 185.49 |
| 26 | Suriname | 31.29 | 77.03 | 12.28 | 0.05 | 503.76 |
| 27 | Sweden | 6.14 | 5.52 | 4.22 | -1.09 | 21.97 |
| 28 | Thailand | 4.4 | 4.97 | 3.07 | -1.13 | 25.08 |
| 29 | Tunisia | 5.47 | 2.62 | 5.1 | 0.28 | 14.2 |
| 30 | United Kingdom | 2.32 | 2.44 | 1.51 | -1.64 | 9.01 |
| 31 | Uruguay | 36.1 | 32.25 | 25.15 | 4.15 | 125.57 |
| 32 | Venezuela | 72.36 | 129.21 | 25.5 | 1.62 | 500.32 |

**Bayesian Model Checks and Evaluations**

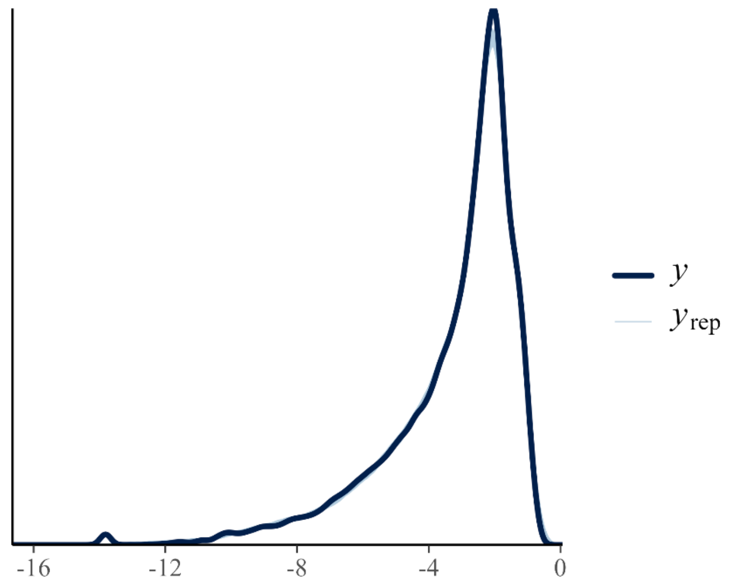


**Figure A1**. Comparison of the density of observed fertility rates with the density of replicated fertility rates. The x-axis shows fertility rates on a logit scale, and the y-axis shows the density of those rates. The dark blue line indicates the density of the observed fertility rates, while each light blue line shows the density of a replicate.

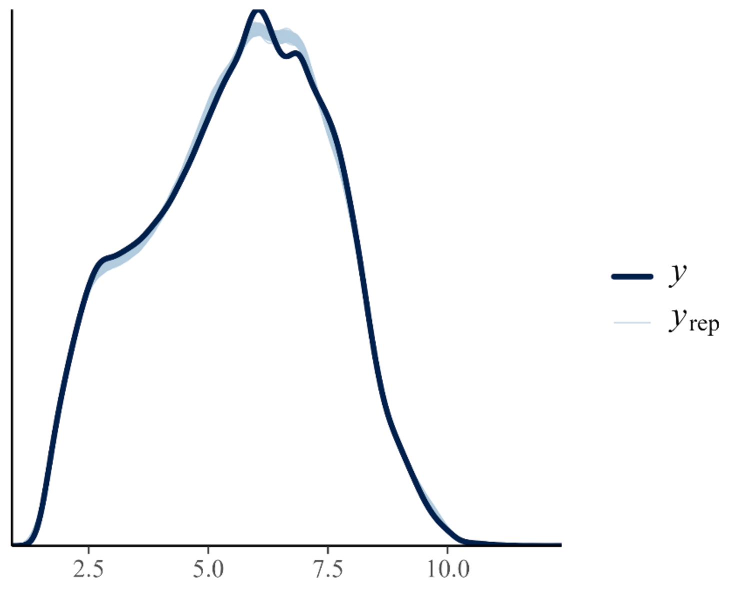

**Figure A2**. Comparison of the density of observed survival ratios with the density of replicated survival ratios. The x-axis shows survival ratios on a logit scale, and the y-axis shows the density of those rates. The dark blue line indicates the density of the observed survival ratios, while each light blue line shows the density of a replicate.

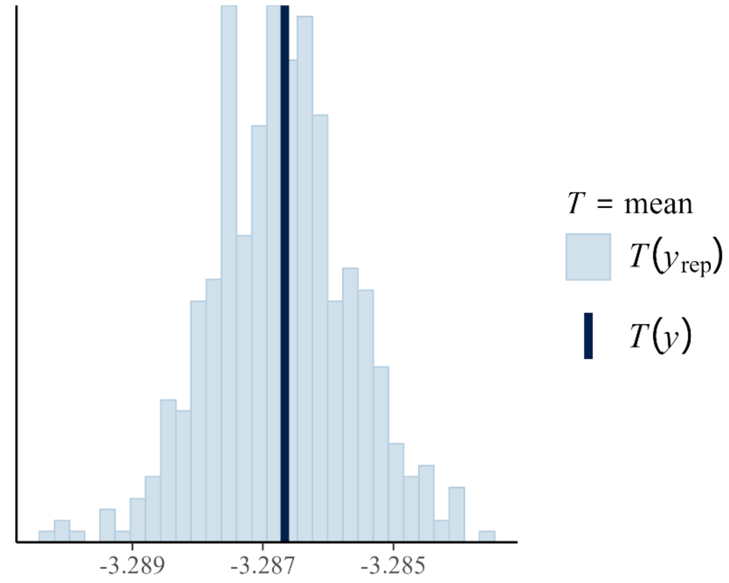


**Figure A3**. Histogram of the test statistic $T[\boldsymbol{y_{rep}}, \boldsymbol{\theta}]$ mean for the replicated fertility rates. The vertical dark blue line, approximately at the centre of the histogram, shows the mean of the observed fertility rates.

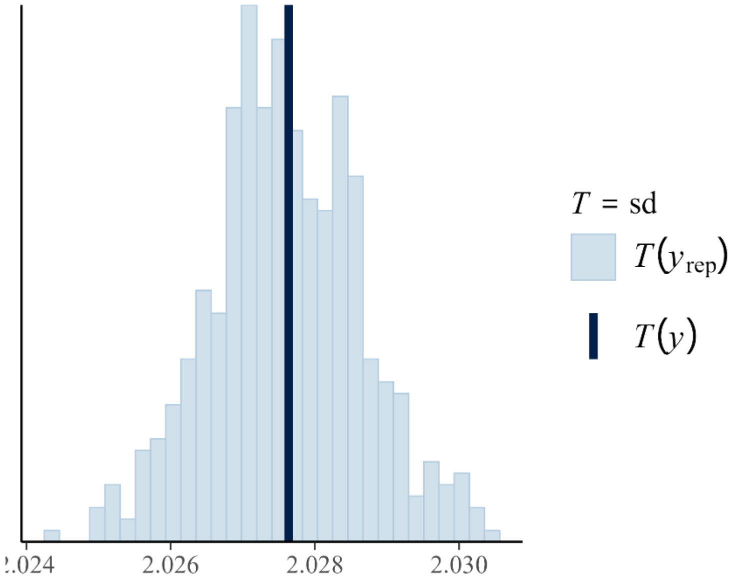

**Figure A4**. Histogram of the test statistic $T[\boldsymbol{y_{rep}}, \boldsymbol{\theta}]$ standard deviation for the replicated fertility rates. The vertical dark blue line, approximately at the centre of the histogram, shows the standard deviation of the observed fertility rates.

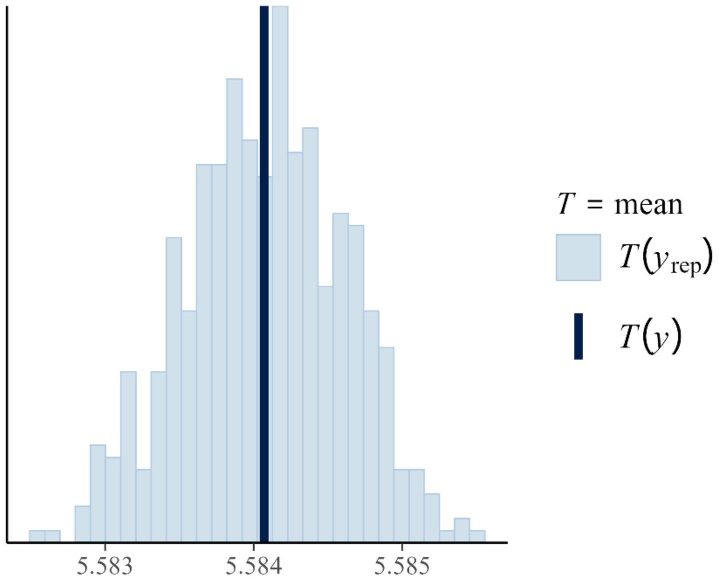


**Figure A5**. Histogram of the test statistic $T[\boldsymbol{y_{rep}}, \boldsymbol{\theta}]$ mean for the replicated survival ratios. The vertical dark blue line, approximately at the centre of the histogram, shows the mean of the observed survival ratios.

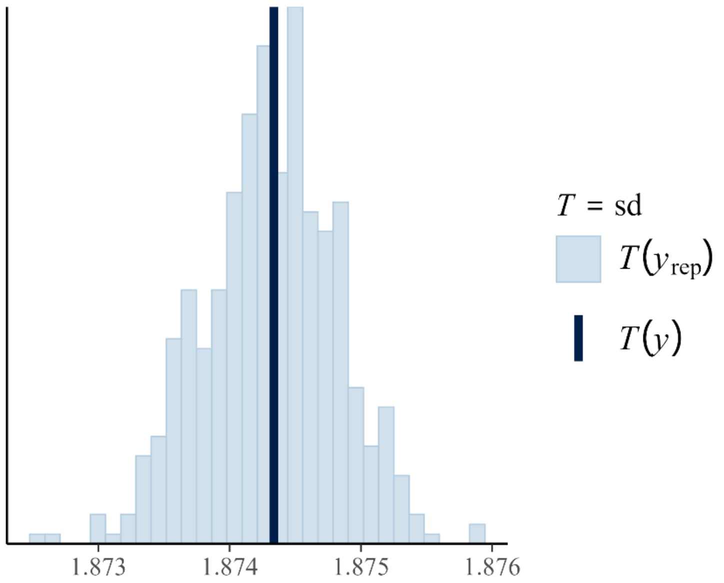

**Figure A6**. Histogram of the test statistic $T[\boldsymbol{y_{rep}}, \boldsymbol{\theta}]$ standard deviation for the replicated survival ratios. The vertical dark blue line, approximately at the centre of the histogram, shows the standard deviation of the observed survival ratios.

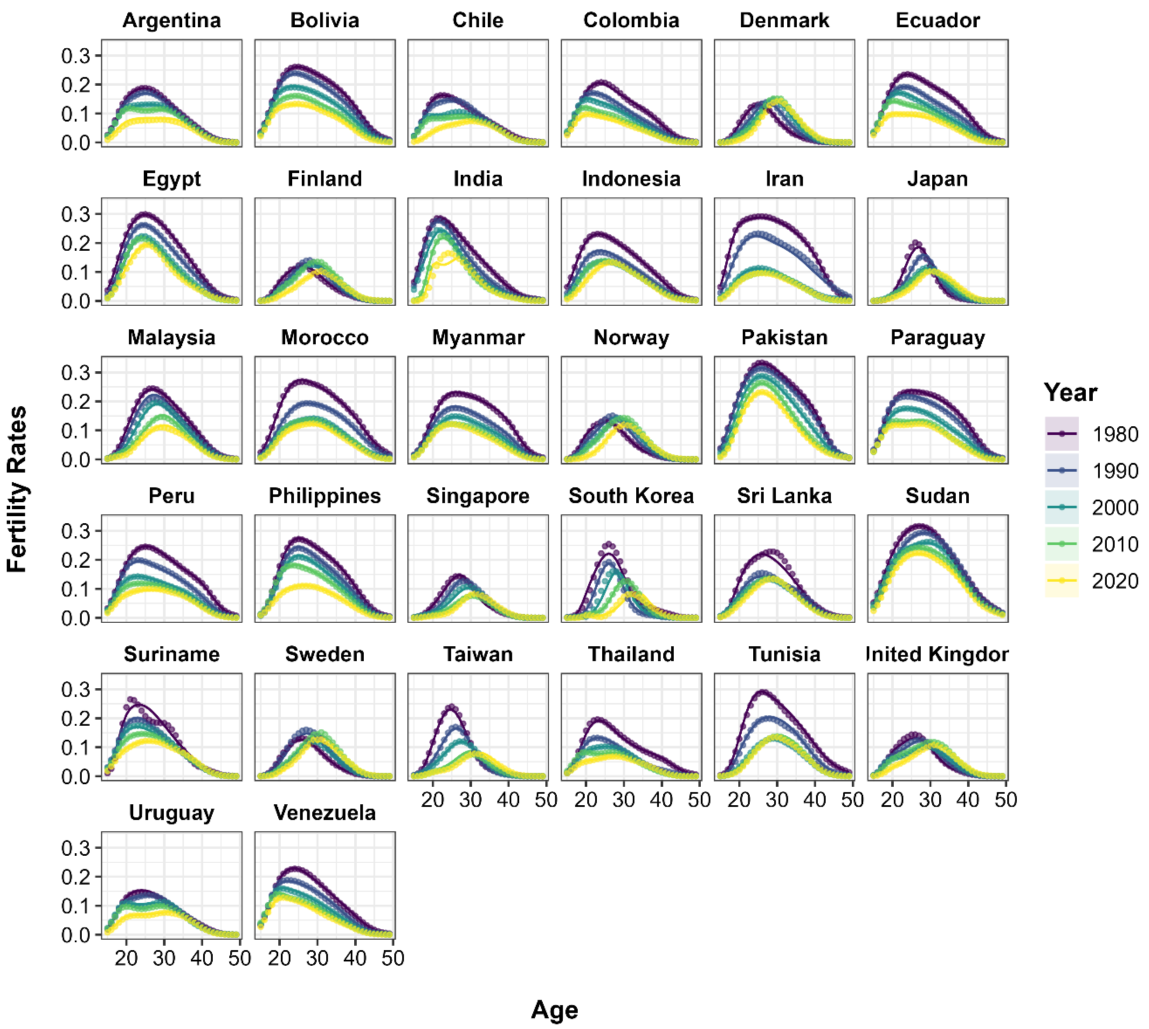


**Figure A7**. Age pattern of observed vs fitted fertility rates for all 32 examined countries. We presented the median of the posterior draws of fertility rates, represented by solid lines, and the 95 per cent uncertainty interval, shown by a band of the same colour. The observed rates are represented by the points. We have presented five select years out of the 51 examined years, and each year is shown in a different colour, ranging from yellow to blue.

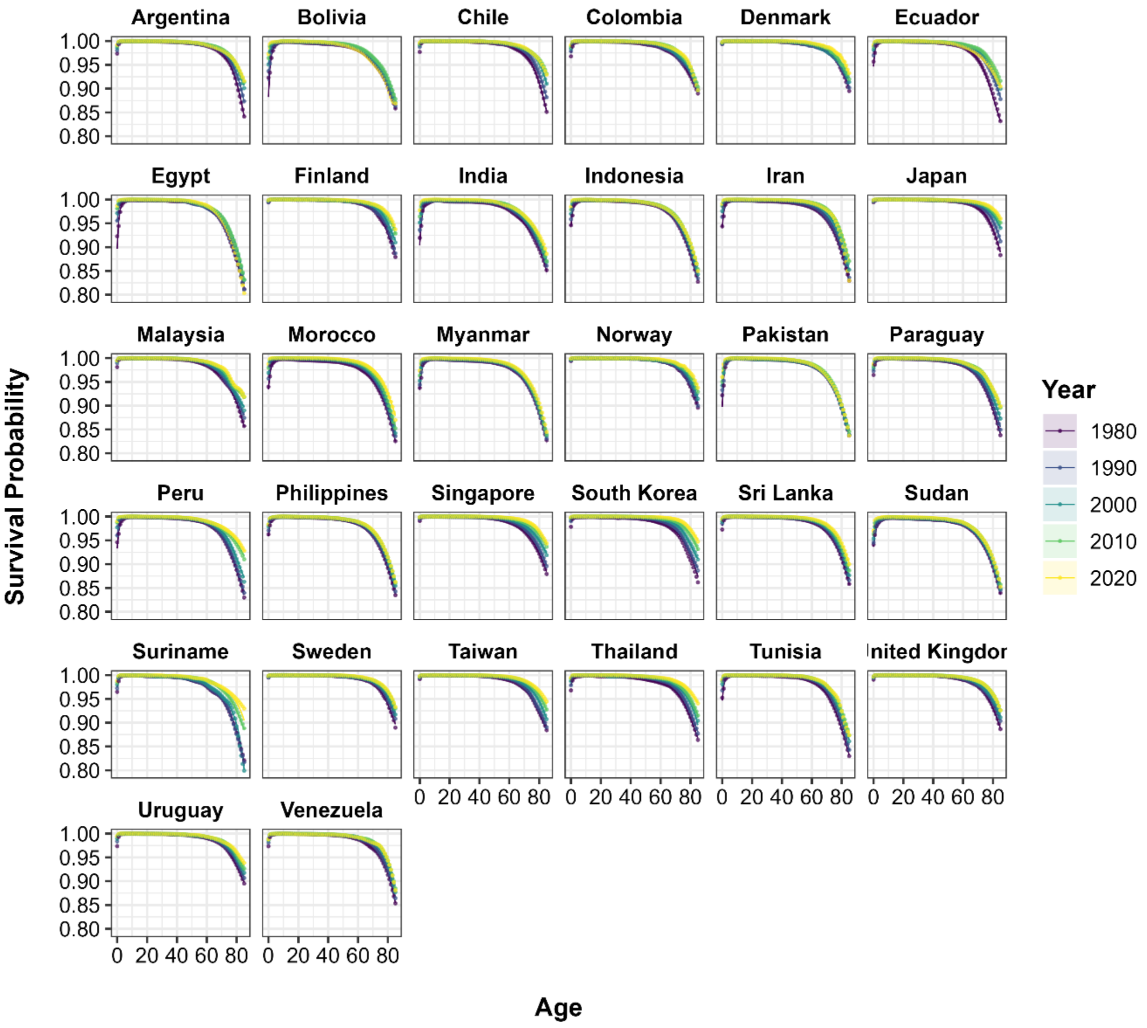


**Figure A8**. Age pattern of observed vs fitted survival ratios for all 32 examined countries. We presented the median of the posterior draws of survival ratios, represented by solid lines, and the 95 per cent uncertainty interval, shown by a band of the same colour. The observed rates are represented by the points. We have presented five select years out of the 51 examined years, and each year is shown in a different colour, ranging from yellow to blue.

**Table 2**. Root mean squared error (RMSE) and mean absolute error (MAE) values of the Bayesian hierarchical model for fertility rates and survival ratios.

| Model | RMSE | MAE |
|---|---|---|
| **Fertility Model** | 0.0037 | 0.0021 |

| Model | RMSE | MAE |
| --- | --- | --- |
| **Survival Model** | 0.0016 | 0.0005 |

#### Additional Figures

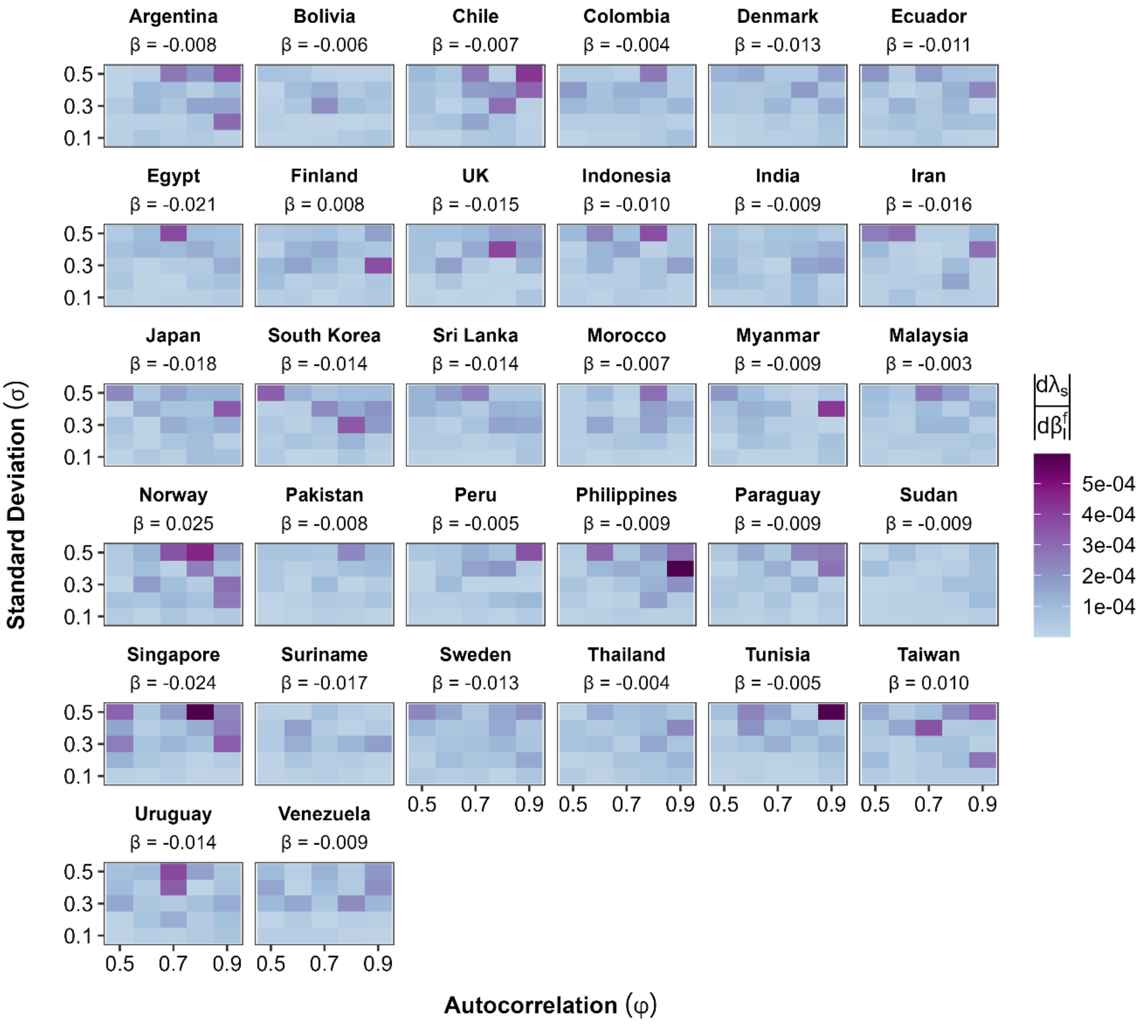


**Figure A9.** The absolute value of the stochastic growth rate's sensitivity to perturbations in the slope coefficient of inflation on fertility rates $\left|\frac{d\lambda_s}{d\beta_i^f}\right|$ is higher at higher variance and autocorrelation in most of the examined countries. Higher absolute values of the stochastic sensitivities are shown in dark purple, and the light blue colours represent the lowest sensitivities. The range of temporal autocorrelation on the x-axis and standard deviation on the y-axis shows the parameter grid used to simulate the inflation time series over 1000 steps. Hence, each element of the matrix plot shows the stochastic sensitivity for one standard deviation-autocorrelation combination.

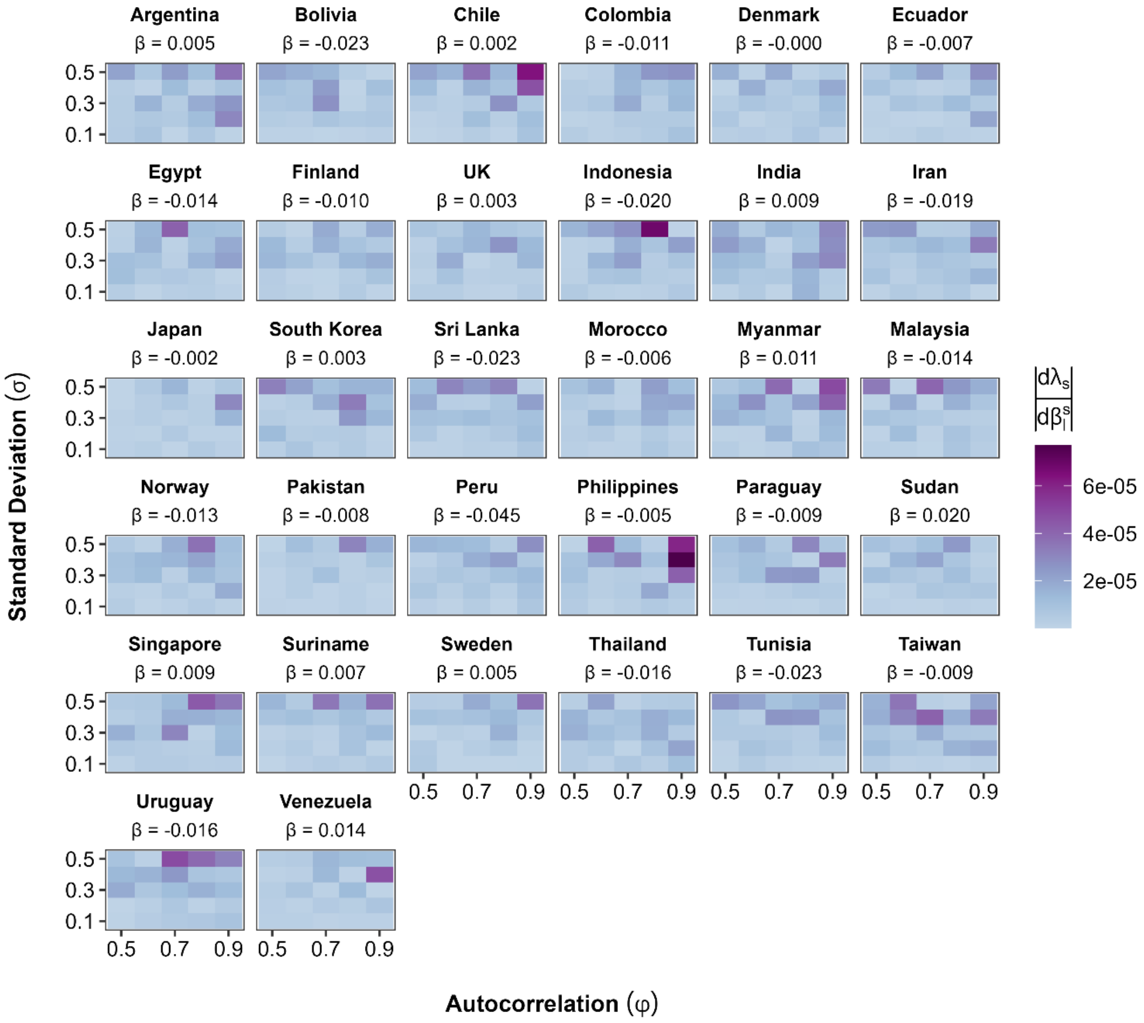


**Figure A10.** The absolute value of the stochastic growth rate's sensitivity to perturbations in the slope coefficient of inflation on survival ratios $\left|\frac{d\lambda_s}{d\beta_i^S}\right|$ is higher at higher variance and autocorrelation in most of the examined countries. Higher absolute values of the stochastic sensitivities are shown in dark purple, and the light blue colours represent the lowest sensitivities. The range of temporal autocorrelation on the x-axis and standard deviation on the y-axis shows the parameter grid used to simulate the inflation time series over 1000 steps.
